# A bifunctional antibody conjugate marks the location of DNA binding proteins on deproteinized DNA fibers

**DOI:** 10.1101/2024.08.29.609705

**Authors:** Durga Pokharel, Althaf Shaik, Himabindu Gali, Chen Ling, Marina A. Bellani, Michael M Seidman

## Abstract

Immunofluorescent foci of DNA Damage Response (DDR) proteins serve as surrogates for DNA damage and are frequently interpreted as denoting specific lesions. For example, Double Strand Breaks (DSBs) are potent inducers of the DDR, whose best-known factor is the phosphorylated histone variant H2AX (γ-H2AX). The association with DSBs is so well established that the reverse interpretation that γ-H2AX invariably implies DSBs is routine. However, this conclusion is inferential and has been challenged. The resolution of this question has been hampered by the lack of methods for distinguishing the location of DDR proteins relative to DSBs caused by sequence indifferent agents. Here, we describe an approach for marking the location of DDR factors in relation to DSBs on DNA fibers. We synthesized a two-arm “Y” conjugate containing biotin and trimethylpsoralen (TMP) coupled to a secondary antibody. After exposure to a DNA breaker, permeabilized mammalian cells were incubated with a primary antibody against the DDR factor followed by binding of the secondary antibody in the conjugate to the primary antibody. Exposure to longwave UV light covalently linked the psoralen to the DNA. DNA fibers were spread, and the immunofluorescence of the biotin tag denoted the location of the target protein.

Graphical abstract

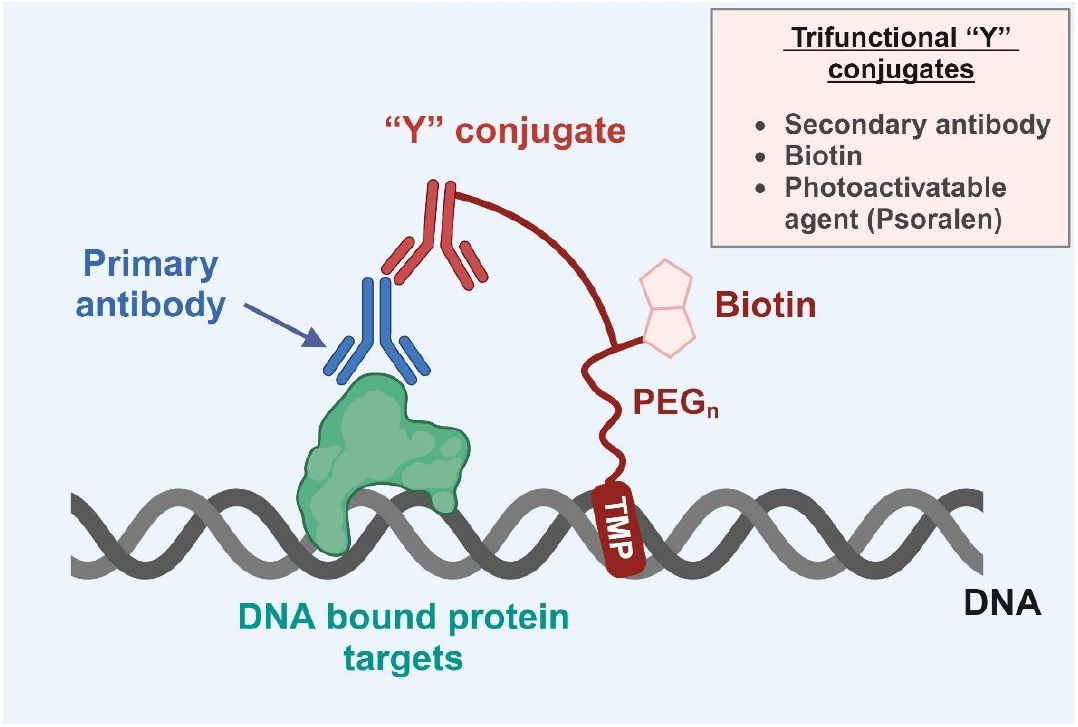

## INTRODUCTION

DNA is under continual assault from both exogenous and endogenous reactive agents, most of which introduce multiple forms of damage. For example, ionizing radiation causes single and double-strand breaks and oxidative base modifications ^1^. The response to DNA damage termed the DDR, involves hundreds if not thousands of proteins that are involved in numerous pathways, among them chromatin remodeling, repair, activation of stress response pathways, and control of replication ^2^. Recruitment of DDR factors to sites of damage can be recognized by immunological assays such as immunofluorescence and chromatin immunoprecipitation. Although these assays are very powerful and widely applied, identification of the inducing lesion is often inferential. For example, if a particular lesion induces the DDR, then it is often assumed that activation of the DDR indicates the presence of that lesion regardless of the multiple forms of damage induced by the DNA reactive agent.

Among the numerous kinds of DNA damage, DSBs are considered the most dangerous. They are induced by a variety of treatments, including ionizing radiation. Unrepaired breaks can provoke genomic rearrangements that initiate tumorigenesis, chromosome breakage, and loss, and the activation of inflammatory and apoptotic pathways. Within seconds to minutes after introduction, they induce the formation of what is perhaps the best-known DDR factor, the histone variant H2AX phosphorylated at serine 139 ^3^. γ-H2AX can be detected with specific antibodies that have been used for many years in immunofluorescence assays to mark sites of DNA damage in fixed cells ^4^. Because the original discovery of γ-H2AX was in the context of DSBs, the appearance of this mark is often reported, ipso facto, as proof of DSBs. However, this interpretation has been questioned, and it has been argued that the association of γ-H2AX is not exclusive to DSBs ^5-8^. Resolution of this issue has been complicated. Attempts to assess the relative frequency of γ-H2AX associated with DSB vs γ-H2AX at non-DSB lesions are hampered by the lack of a straightforward technology for these determinations. Immunofluorescent foci in nuclei report the localization of responding proteins without identifying the underlying lesion. On the other hand, electrophoretic demonstrations of DNA breakage, such as the comet assay, are performed under conditions of deproteinization, thus losing the connection between the lesion and DDR factors ^9^.

Methods for spreading deproteinized single DNA molecules from mammalian cells on glass slides have been available for many years ^10, 11^, and are commonly used for studies of DNA replication. These involve short incubations of proliferating cells with nucleoside analogues, such as CldU or BrdU before harvest ^12^. The tracts of incorporated analogues displayed by immunofluorescence report features of DNA replication such as fork speed, fork restart following treatments that stall replisomes, etc.

However, DNA fiber technology enables other, largely unexploited, experimental applications. For example, extended incubations with analogues label DNA uniformly so that very long fibers can be visualized ^13, 14^. Features intrinsic to deproteinized DNA, unrelated to replication, can be detected on these fibers. Notably, and relevant to this report, DNA ends arising from DSBs are apparent ^15^. A method to mark the location of chromatin proteins on fibers would enable a more direct assessment of the connection between a DDR protein such as γ-H2AX and ends and might reveal associations at other, non DSB, sites. In this report, we describe our development of a method to mark the location of a DDR protein on DNA fibers isolated from irradiated cells.

## RESULTS AND DISCUSSION

Our protocol to mark the location of a chromatin protein on DNA fibers involves incubation of permeabilized cells with a primary antibody against a DDR protein, followed by binding of the primary with a secondary antibody linked to both a visualization tag and also to a reagent whose covalent reaction with DNA is under investigator control (**Fig 1a**). The conjugate contained three elements

**Figure 1.**
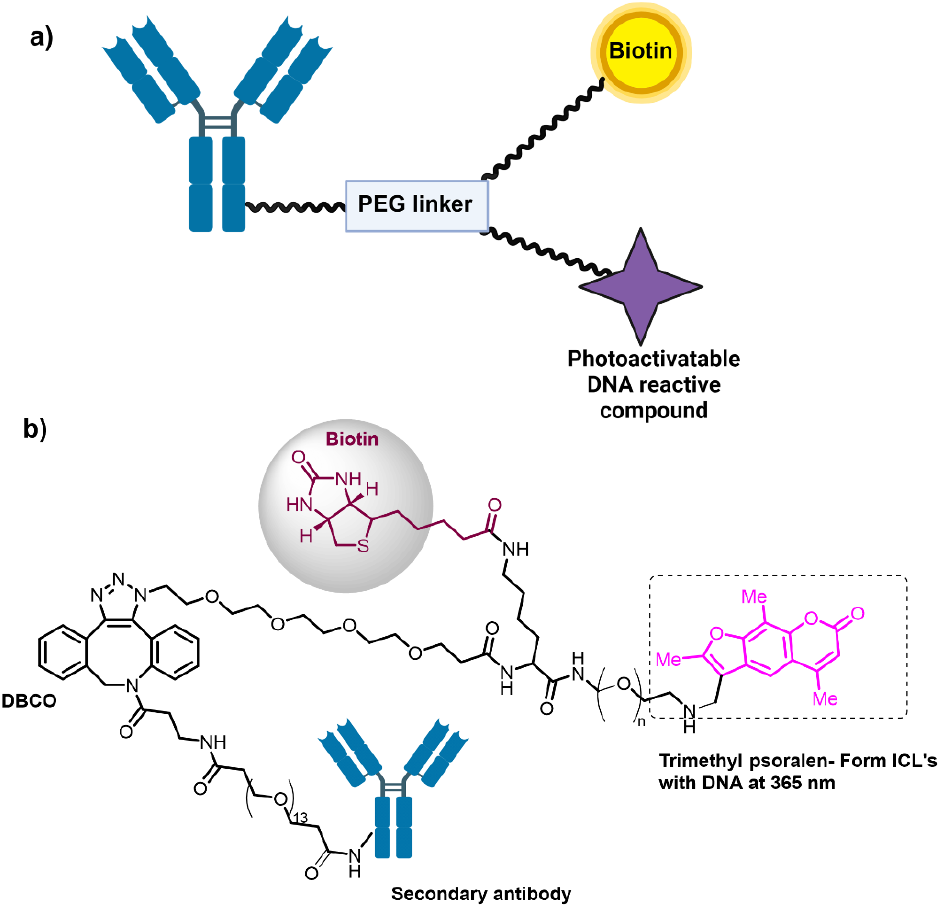
**a)** Design strategy of an antibody-biotin-psoralen Y conjugate for marking the location of chromatin proteins on DNA fibers; **b)** chemical structure of Y conjugate.

1. A component for protein recognition. Primary antibodies afford target recognition with high specificity. Although we could have attached the tag and the covalent reactant to primary antibodies, we chose instead to couple them to anti-mouse and anti-rabbit secondary antibodies. Thus, they could be employed in conjunction with primary antibodies of either species eliminating the need for synthesis of a reagent for each target.
2. An imaging tag: We used biotin, frequently used for capture and visualization.
3. An investigator-controlled DNA reactive moiety-Psoralens are DNA intercalators that are reactive only during exposure to long-wave ultraviolet light (UVA). They form interstrand crosslinks (ICLs) between diagonal thymine bases at 5’TA: AT sites and can accept side chains without loss of crosslinking activity. The requirement for photoactivation allows for binding of the 2° antibody conjugate followed by stringent washing to remove unbound reagent before the reaction with DNA. The experimental procedure involves long-term incubation of proliferating cells with nucleoside analogues such as CldU, permeabilization of cells, incubation of nuclei with a target-specific primary antibody, removal of the unbound antibody, incubation with the conjugate (hereafter referred to as the Y conjugate), removal of the unbound conjugate, exposure of the nuclei to UVA, nuclear disruption and chromatin deproteinization by suspension in sodium dodecyl sulfate, spreading DNA fibers, display of fibers by immunofluorescence against incorporated nucleoside analogues and an immuno quantum dot for the biotin tag.

### Synthesis of Y conjugates

We synthesized three versions of the Y conjugate. Each contained a stem terminating with an azide and had varying lengths of the linker (n= 10, 15, 23) between the fork in the Y and the trimethyl psoralen (**Fig 1b**) (**Scheme 1**). Anti-mouse and anti-rabbit secondary antibodies were reacted with an NHS ester linked to dibenzyloctyne (DBCO) ^16^ by a PEG13 linker **(Scheme 2**). The DBCO-derivatized secondary antibodies were then conjugated to the Y compounds by azide: DBCO coupling to form either an anti-rabbit or an anti-mouse Y conjugate **(Scheme 3)**. We performed a series of control experiments to assess the activity of the three components of the Y conjugate. The integrity of the biotin on the conjugate was determined by denaturing SDS polyacrylamide gel electrophoresis followed by Western blot analysis of the biotin by streptavidin coupled to peroxidase (**Fig 2**) ^17, 18^. This revealed undegraded bands of 25 and 50 K MW as expected for heavy and light antibody chains. Consequently, both biotin and antibody were intact.

**Schemes 1.**
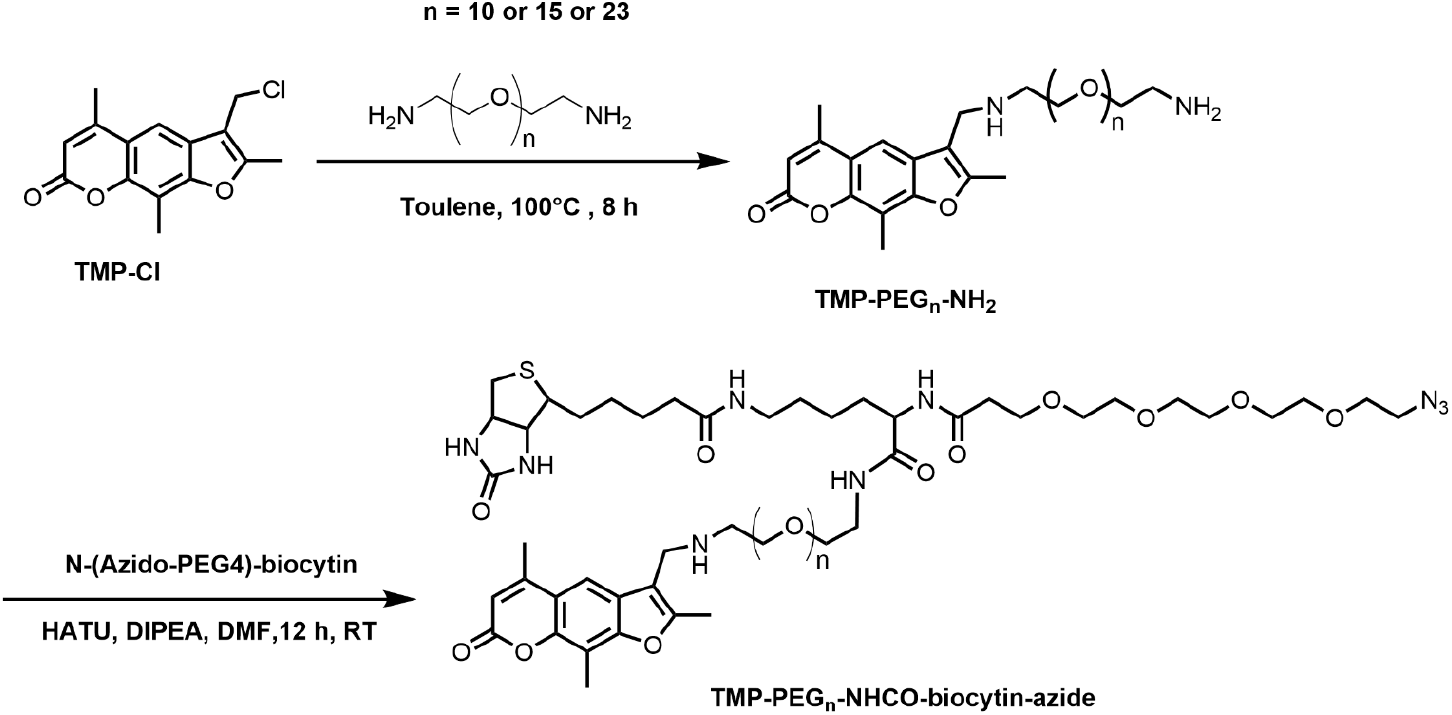
Synthesis of TMP-PEG_n_-biocytin azide containing (n = 10 or 15 or 23) clickable handles.

**Scheme 2.**
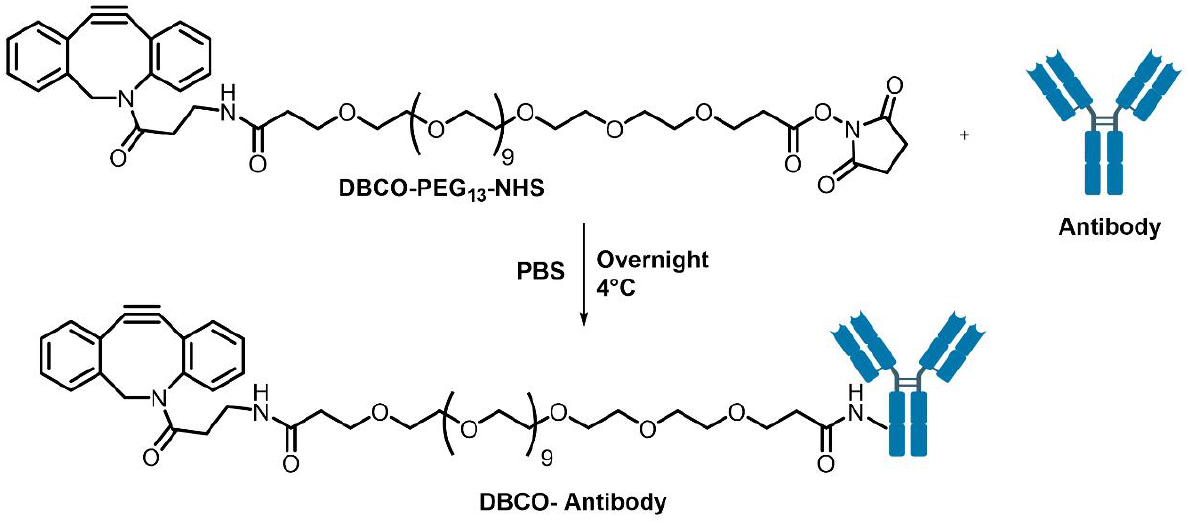
Strained alkyne installed onto the antibody using N-hydroxysuccinimide chemistry

**Scheme 3.**
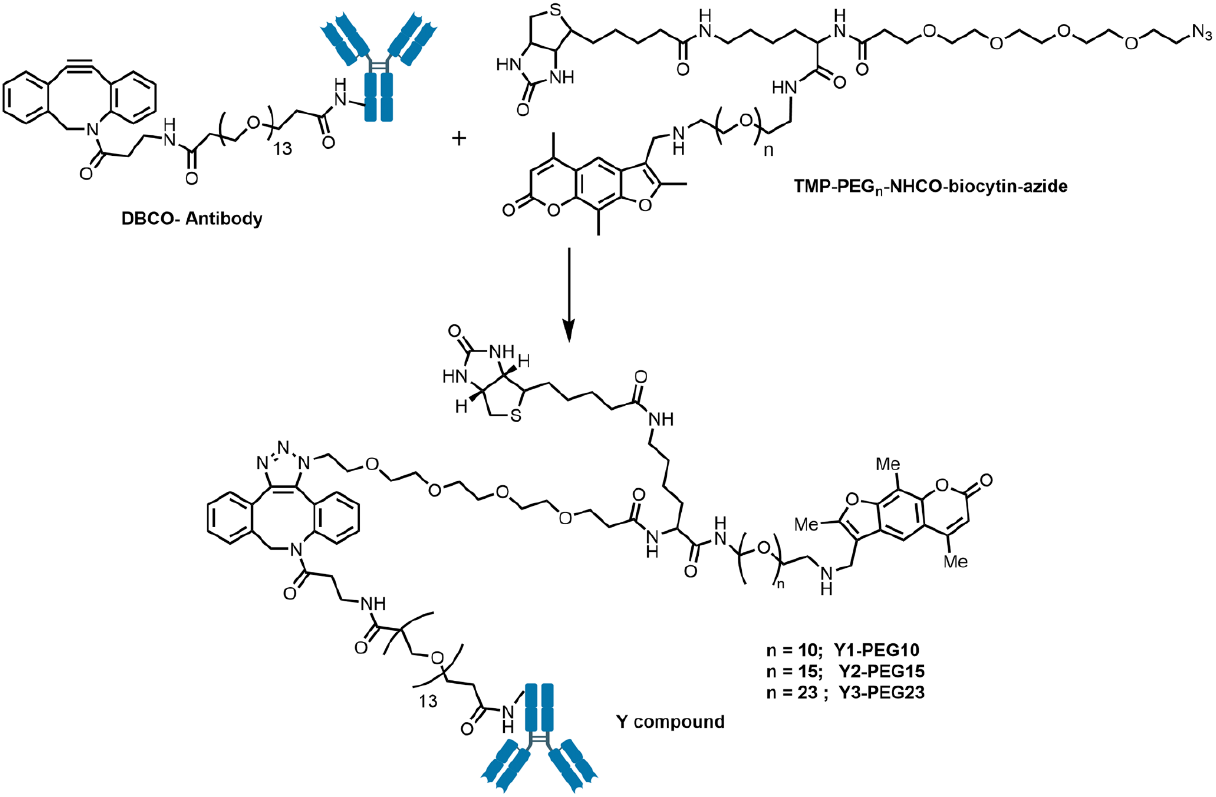
Clickable intermediates were further conjugated to an antibody via Strain-Promoted Azide−Alkyne Cycloaddition (SPAAC) Click Chemistry.

**Figure 2.**
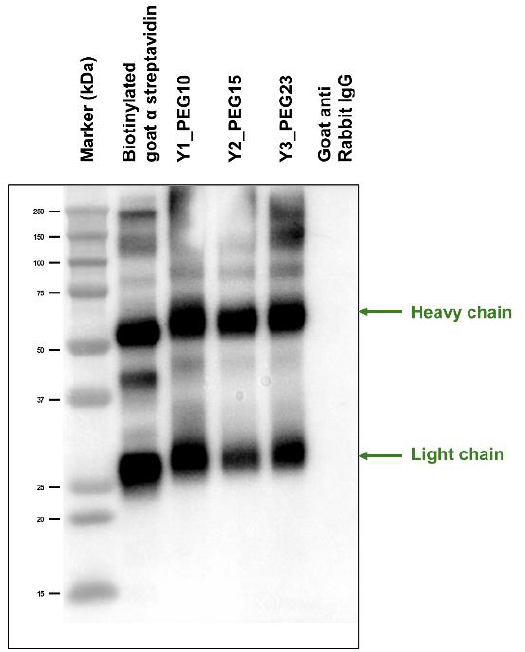
Western blot detection of biotinylated antibody-TMP conjugates (Y conjugates) showing biotin modification. Non biotinylated goat anti rabbit IgG was loaded in the last lane.

The activity of the psoralen was determined by incubation of permeabilized nuclei with the conjugate followed by exposure to UVA, and then extensive washing to remove unreacted reagent. In control experiments, the cells were incubated but the UVA exposure was omitted. The bound reagent was displayed by immunostaining against biotin. Cells treated with both reagent and UVA showed strong nuclear fluorescence, while the control showed no signal (**Fig 3**). This demonstrated the activity of both the psoralen and the biotin. The activity of the coupled secondary antibody in the conjugate was tested in two experiments. In one, cells were treated with Digoxigenin-trimethyl psoralen (Dig-TMP)/UVA to covalently attach the Dig antigen to cellular DNA ^19^. The cells were fixed and incubated with a primary antibody against the Dig tag, and then with either a commercial fluorescent secondary antibody or the different versions of the Y conjugate. The biotin on the conjugates was detected by streptavidin linked to an Alexa fluor (**Fig 4**). In a second experiment, PCNA was the target of the primary antibody followed by a commercial secondary or the Y conjugates (**Fig 5**). These results confirmed the activity of the secondary antibody arm of the conjugates and this experimental series demonstrated that all the components of the Y compounds were functional. A reagent designed to detect and mark sites of specific chromatin proteins should not mark sites in DNA that are not intended targets. The compaction of DNA in nuclear chromatin, and a wealth of chromosome conformation capture experiments, indicate that a site on DNA associated with a target of interest will be near sequences that do not have that target ^20^.

**Figure 3.**
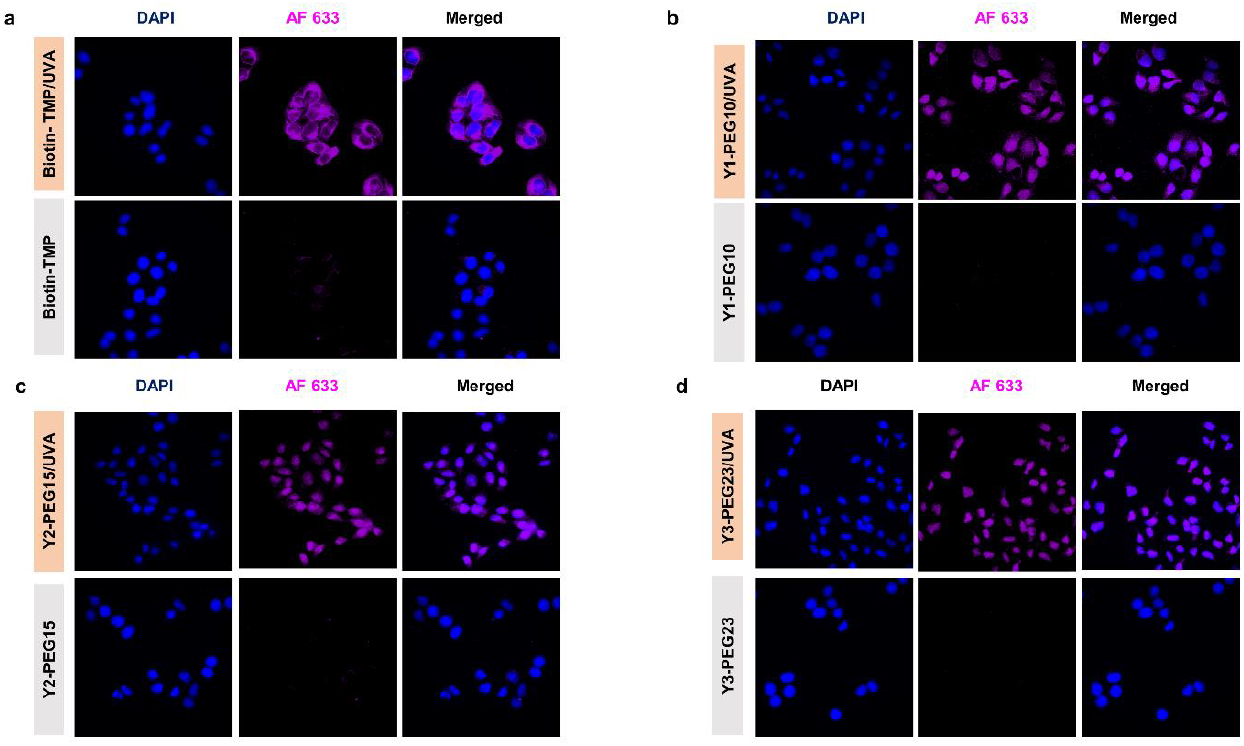
The activity of the psoralen and biotin in the Y conjugates. Permeabilized Hela cell nuclei were incubated with respective Y conjugates and exposed to UVA+/-. The immunofluorescence of the biotin tag is shown in magenta (top panel). Nuclei stained with DAPI are shown in blue. Without UVA treatment no signal was detected in the control

**Figure 4.**
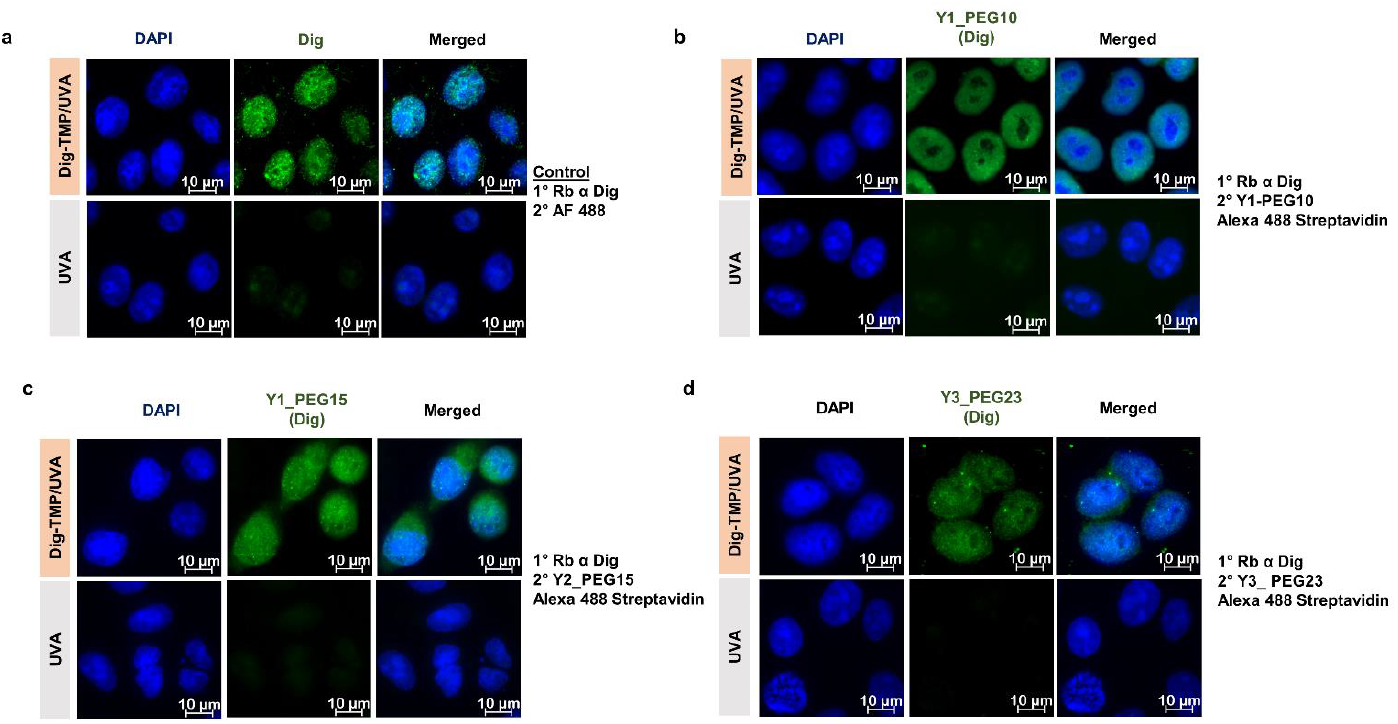
The activity of the antibodies in the Y conjugates. Immunofluorescence detection of the Dig antigen tag in genomic DNA using Y conjugates. Cells were treated with 1 µM Dig-TMP for 20 min, then irradiated with UVA (3 J/cm^2^). The anti-Dig rabbit primary antibody and anti-rabbit Y conjugate detection are shown by the Alexa 488 signal (green) and the nuclear stain by DAPI (blue).

**Figure 5.**
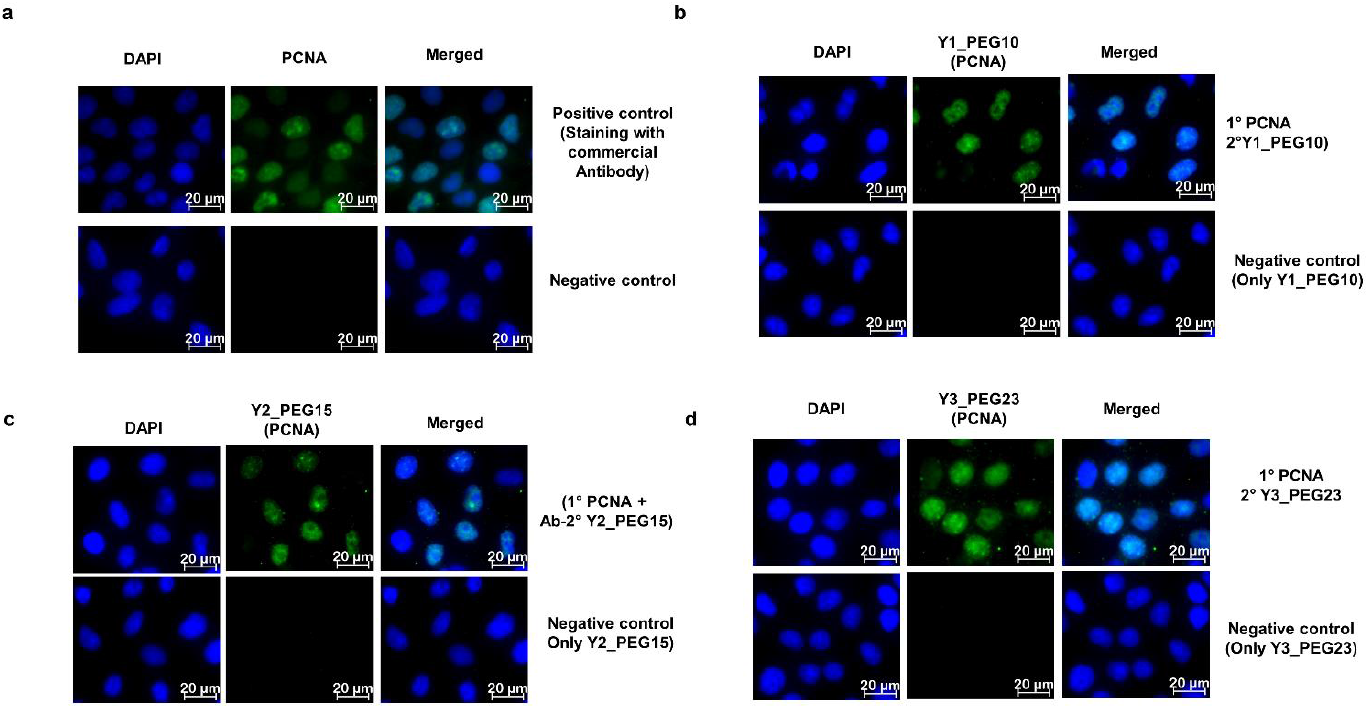
The immunofluorescence detection of PCNA signals in Hela cells using modified secondary Y conjugates against primary PCNA antibodies is shown in green. Nuclei stained with DAPI are shown in blue.

As described above the Y conjugates were constructed with different lengths of the linkers connecting the psoralen. We anticipated that a too-short linker would fail to mark target sites, while one too-long would reach out and mark DNA without the target. Cells were incubated for 24 hrs with CldU and then treated with Dig-TMP/UVA. This introduced a covalent Dig tag in the DNA. The cells were permeabilized, fixed, and then incubated with a primary antibody against Dig. After washing to remove unbound primary antibodies they were incubated with the conjugates of differing linker lengths. After washing to remove unbound conjugates they were exposed to UVA to covalently attach the bound conjugates to DNA. The DNA was spread on glass slides and the CldU was displayed by immunofluorescence. The Dig and the biotin were bound by primary antibodies and secondaries linked to quantum dots of different wavelengths-Q655 for Dig, Q705 for biotin ^13, 21^ (**Fig 6a**). In preliminary experiments, a conjugate with a PEG linker length of 1 did not give signals on the fibers and was not characterized further. However, biotin was detected on fibers in experiments with the Y10, 15, and Y23 versions. The frequencies of colocalized biotin and Dig Q dots and that of the biotin alone were determined for the different constructs (**Fig 6b**). A high frequency of colocalization was observed for Y10, but there was a pronounced reduction in the relative frequency of colocalized and a marked increase in biotin-only signals with the Y15 and 23 constructs. This indicated that the linker arms on the Y15 and Y23 conjugates were too long resulting in marking DNA at nontarget sites. We concluded that the Y10 version was optimal. We then addressed the localization of a DDR marker and DNA breaks and non-break sites. γ-radiation introduces DSBs and other lesions into DNA ^22^, and activates the DDR including the appearance of γ-H2AX foci. Consequently, it was an appropriate DNA damaging agent to examine the relationship between γ-H2AX and the ends of DNA fibers. Cells were incubated with CldU for 24 hrs and then exposed to γ-radiation. This stimulated the appearance of γ-H2AX foci in nuclei as shown by immuno-fluorescence (**Fig 7**). The cells were permeabilized fixed and incubated with a primary antibody against γ-H2AX. They were incubated with the Y10 conjugate and exposed to UVA. Fibers were spread and immunostained to display CldU and biotin. No biotin signals were observed in fiber spreads for cells without radiation exposure, from nuclei in which the primary antibody was omitted, or if there was no UVA after incubation with Y10 (**Fig 8a**). However, in nuclei from irradiated cells in which Y10 incubation was followed by UVA there were biotin signals at DNA ends, but also at internal sites in the fibers. Quantitation of the signals in the two locations revealed that the signals within fibers were more frequent than those at ends (**Fig 8b**). These results demonstrated that following treatment of cells with a DNA damaging agent that introduces multiple lesions, including DSBs, γ-H2AX appears at both DSBs and non-DSB sites, with a greater presence at the latter.

**Figure 6.**
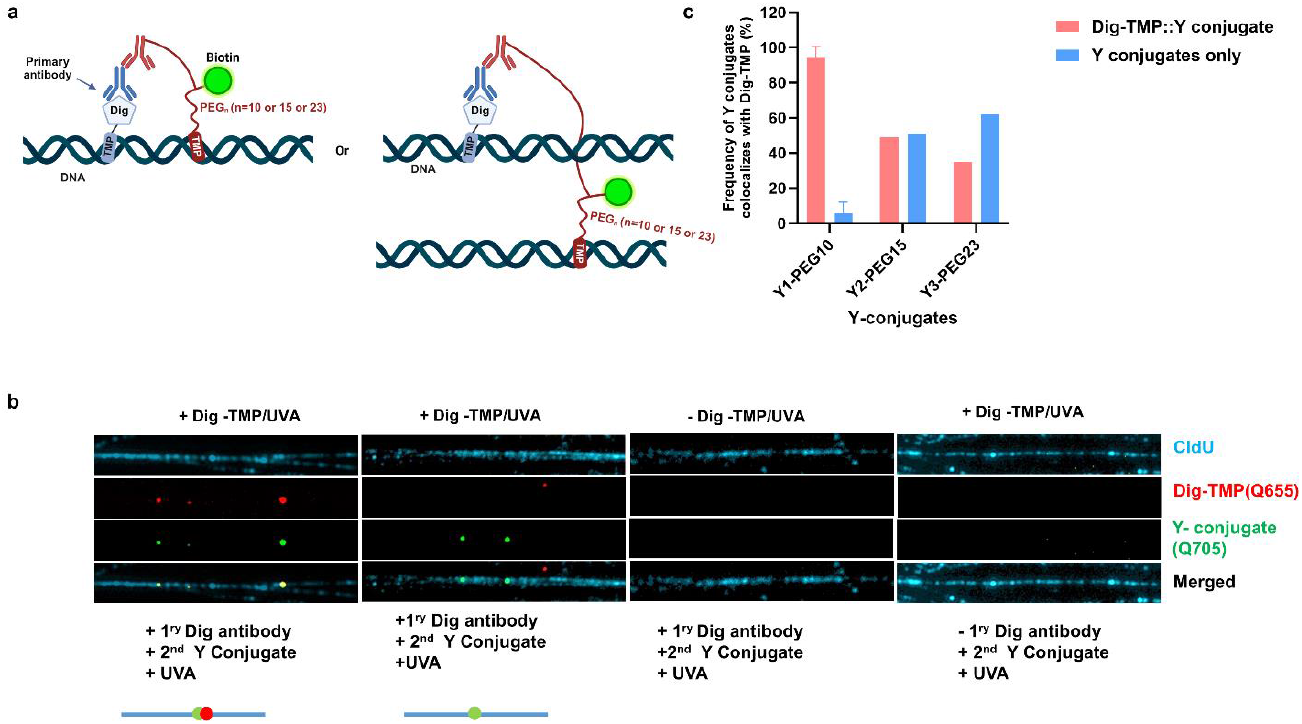
**a)** Schematic diagram showing Y conjugates recognizing Dig TMP ICL on DNA: interpretation of colocalization patterns **b)** DNA fibers assay for various PEG_n_ linkers lengths of Y conjugates to determine the frequencies of colocalized biotin and Dig Q dots. Hela cells were pulsed with CldU (10 µM, 12 h) and then treated with Dig TMP/UVA (1 µM for 20 min, irradiated with UVA 3J/cm^2^). Permeabilized nuclei were incubated with antibodies against Dig and then with respective Y secondary biotin conjugates then UVA 3J/cm^2^. DNA labeled with CldU is shown in light blue, the Dig-TMP signal in red, and biotin in green. From right to left the panels show 1. a fiber with coincident Dig and biotin signals demonstrating the accurate marking on the fiber of the target (Dig) by the biotin in the Y conjugate; 2. A fiber with biotin but no Dig, denoting marking by the Y conjugate on a fiber adjacent to one with the Dig target; 3. Fiber from cells not exposed to Dig-TMP has no biotin signal indicating no nonspecific binding by the primary antibody and Y conjugate secondary; 4. Fibers from cells exposed to Dig-TMP but without incubation with the primary anti-Dig antibody do not have a biotin signal, indicating no nonspecific binding by the Y conjugate. c) Quantification of frequencies of colocalized biotin and Dig Q dots on DNA fiber with different Y conjugates. After the first analysis, the Y2-PEG15 and Y3-PEG23 showed extensive off-target activity and were not further characterized.

**Figure 7.**
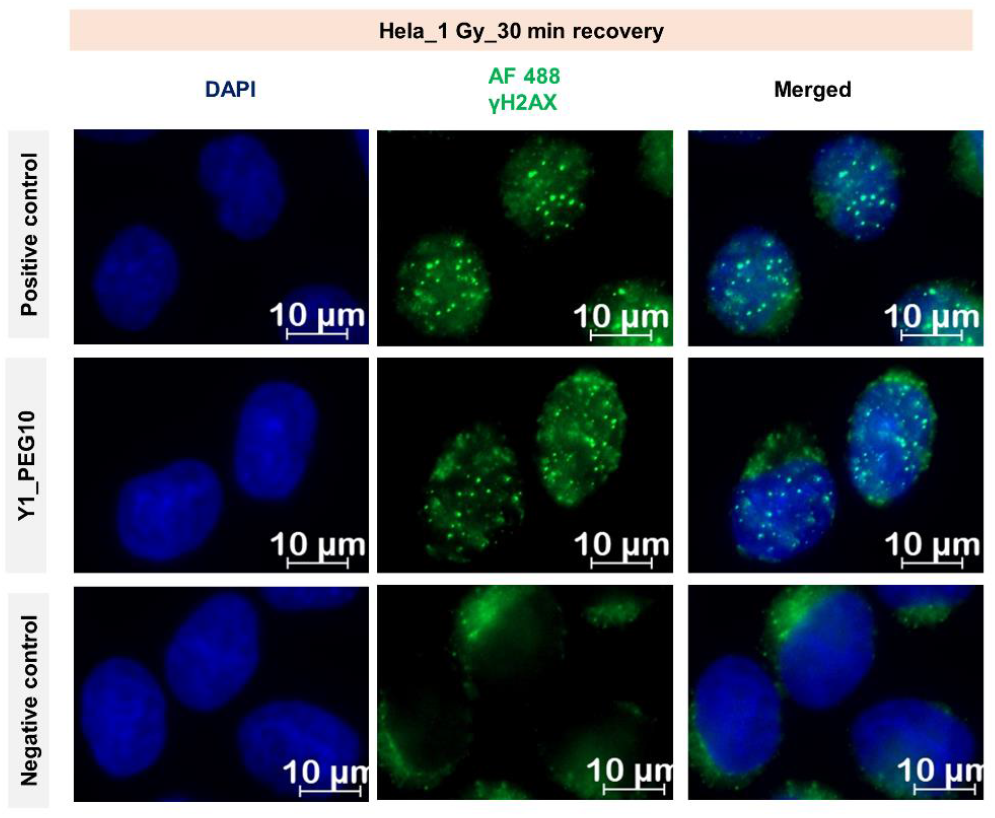
Immunofluorescence detection of γ-H2AX signal using modified secondary Y1-PEG10 conjugate. Hela cells were irradiated with IR (1GY) and then incubated at 37°C for 30 min. Permeabilized cells were blocked and incubated with anti γ-H2AX antibody and then the Y1 PEG10 conjugate. Biotin was visualized by fluorescent streptavidin in a green channel, and Nuclei stained with DAPI shown in blue.

**Figure 8.**
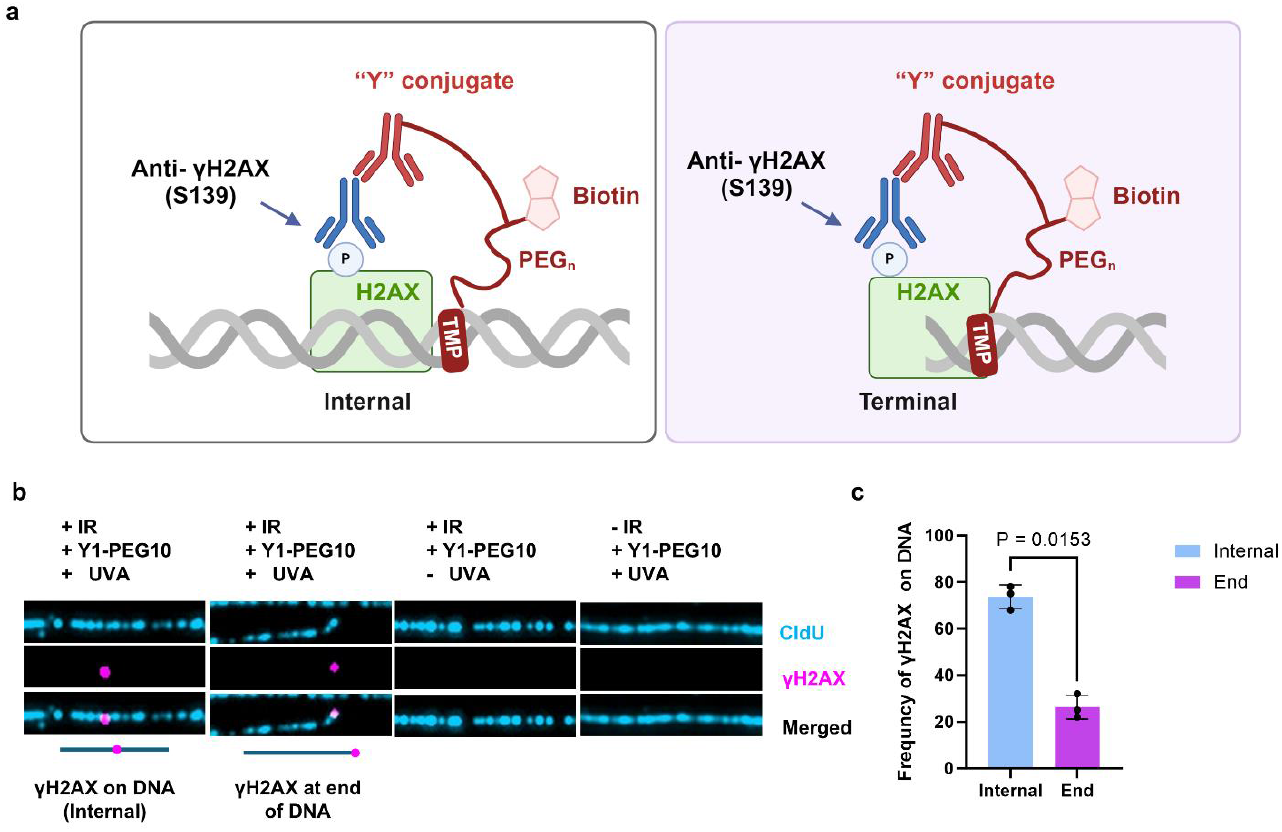
**a)** Depiction of marking γ-H2AX on DNA fiber **b)** Frequency of γ-H2AX foci on DNA. Cells were incubated with CldU (10 µM, 12 h and then treated with IR (1GY) and incubated at 37°C for 30 min. Permeabilized nuclei were incubated with a primary antibody against γ-H2AX and then with the Y1-PEG10 conjugate, then exposed to UVA. DNA labeled with CldU is shown in cyan, and γ-H2AX marked by biotin is shown in magenta; **c)** Quantification of the frequency of γ-H2AX signal at ends and internal sites on DNA fibers.

The strategy described in this report takes advantage of a non-conventional application of DNA fiber methodology. Our results agree with prior proposals that γ-H2AX foci are not uniquely at DSBs ^6-8^ and enable a quantitative assessment of the distribution of this marker at DSB and non-DSB sites. This calculation has practical implications. DSBs can be deadly and agents that induce them, such as radiation, are frequently used in cancer chemotherapy. There is a need for effective measures of the toxic lesions in DNA resulting from exposure to these drugs. Nuclear immunofluorescence of γ-H2AX has been discussed as a diagnostic of DNA DSBs in clinical samples recovered after treatment ^23, 24^. However, we suggest that the fiber assay described here reports a more accurate interpretation of the relationship between γ-H2AX and DSBs.

## AUTHOR INFORMATION

## Present Addresses

Durga Pokharel, Horizon Discovery, Lafayette, CO 80026, USA

Althaf Shaik, Laboratory of Molecular Biology and Immunology, National Institute on Aging, National Institutes of Health, Baltimore MD 21224.

Himabindu Gali, Frederick National Laboratory for Cancer Research, Frederick, MD 21703, USA

Chen Ling, Laboratory of Molecular Biology and Immunology, National Institute on Aging, National Institutes of Health, Baltimore MD 21224.

Marina A Bellani, Laboratory of Molecular Biology and Immunology, National Institute on Aging, National Institutes of Health, Balti-more MD 21224.

## Author Contributions

DP, AS, HG, CL, MAB generated reagents and performed experiments. AS, MAB, MMS wrote the manuscript, MMS conceived the project and designed the experiments.

## Notes

The authors declare no competing financial interests.

## ACKNOWLEDGMENT

This research was supported by the Intramural Research Program of the NIH, National Institute on Aging, United States (Z01-AG000746-08).

## Notes

### Competing Interest Statement

The authors have declared no competing interest.

