## Supplemental data for "A bifunctional antibody conjugate marks the location of DNA binding proteins on deproteinized DNA fibers"

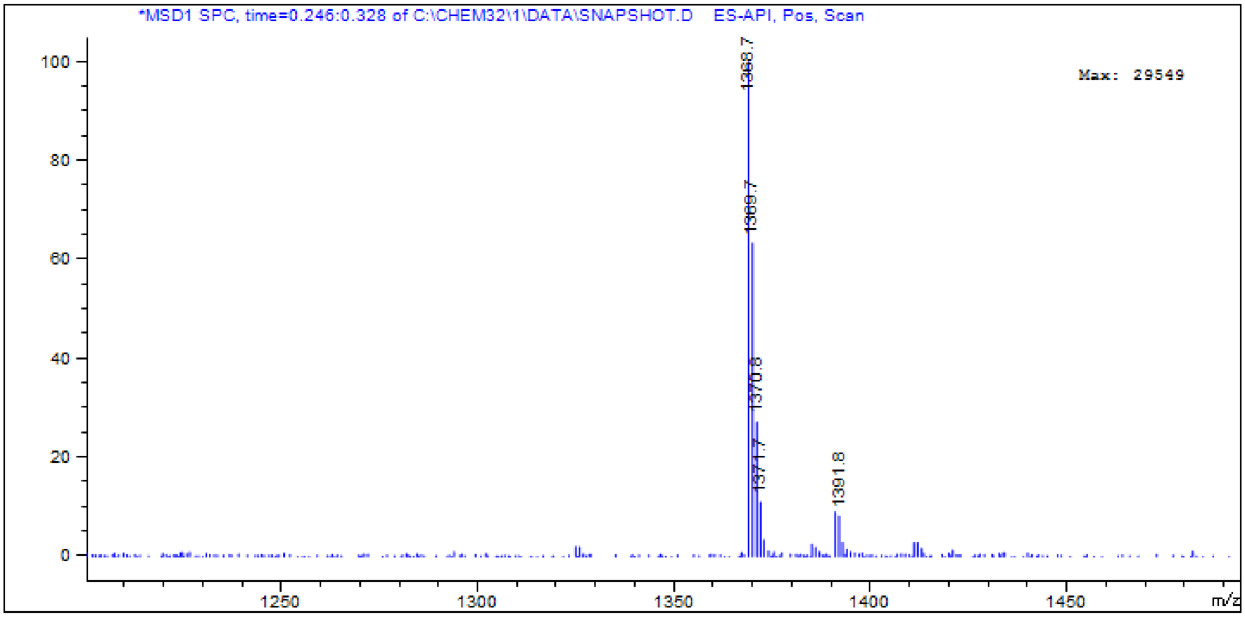


**Figure S1**: LC-MS data for TMP-PEG10-NHCO-Biocytin-azide

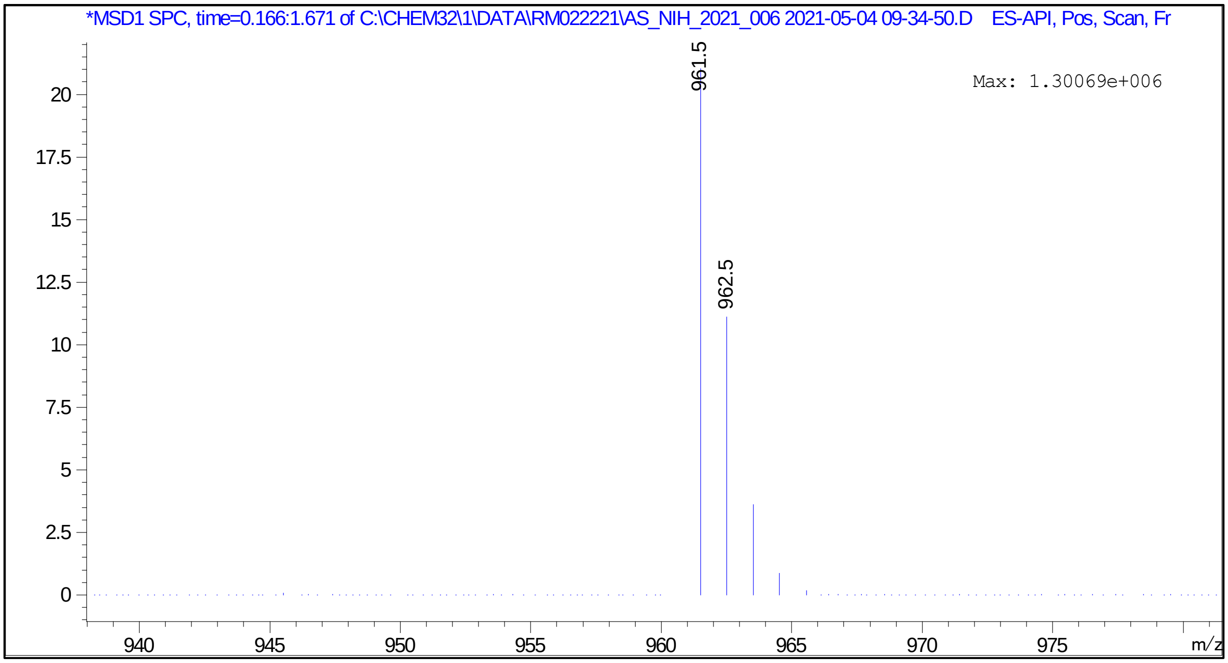


**Figure S2:** LC-MS data for TMP-PEG15-NH2

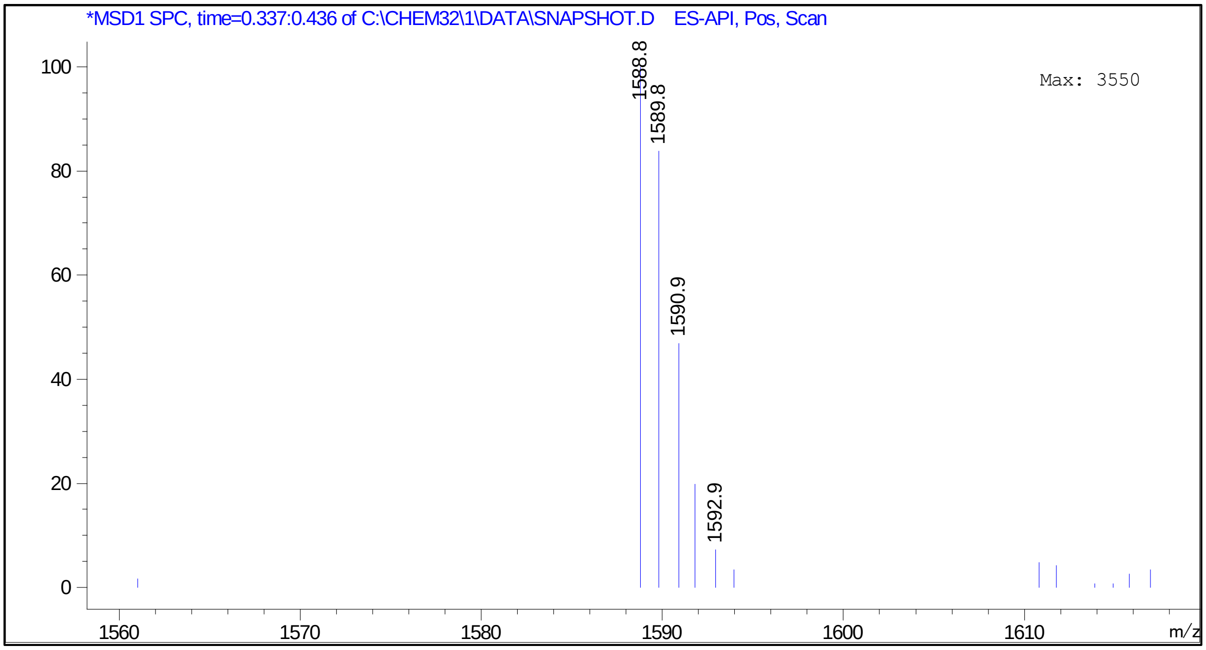


**Figure S3**: LC-MS data for TMP-PEG15-NHCO-Biocytin-azide

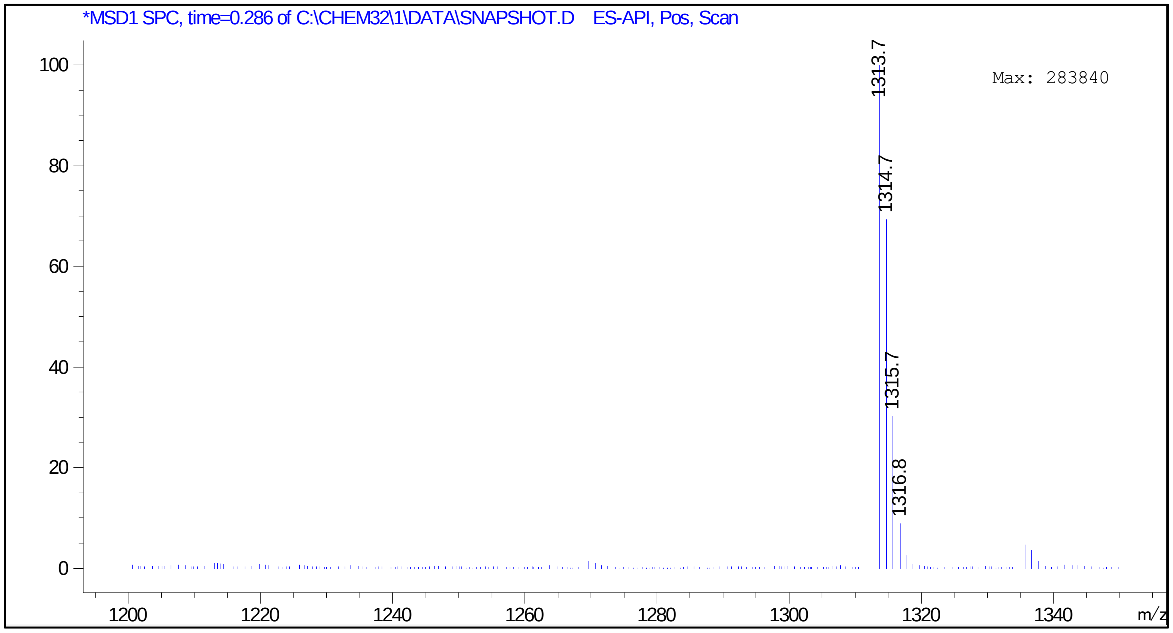


**Figure S4**: LC-MS data for TMP-PEG23-NH2

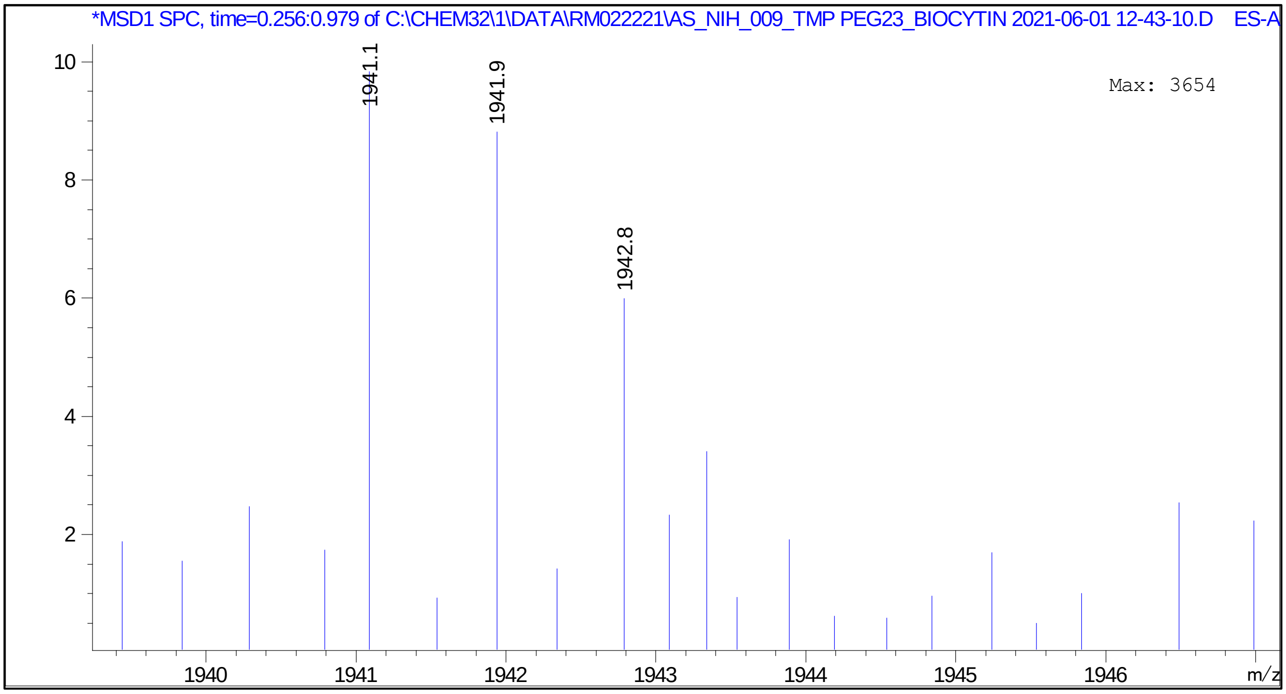


**Figure S5:** LC-MS data for TMP-PEG23-NHCO-Biocytin azide

### Materials and Methods

Starting materials, solvents, and reagents were procured from Sigma Aldrich, Click Chemistry Tools, Broad Pharma, and Roche Chemicals. Reactions were monitored by liquid chromatography-mass spectroscopy (LCMS) or thin-layer chromatography (TLC).

**Synthesis of TMP-PEG$_n$-NH$_2$:** (n= 10, 15, or 23) To the solution of 4′-chloromethyl-4,5′,8-trimethylpsoralen (30 mg, 0.108 mmol) in toluene was added corresponding PEG$_n$ diamines (n= 10 or 15 or 23) (1.086 mmol) and heated at 100 °C for 8 h. The excess solvent was removed under reduced pressure and the product was subjected to silica gel column chromatography and eluted the compound in chloroform, methanol, and 28% ammonia (9:1:0.5).

**TMP-PEG$_{10}$-NH$_2$**: sticky solid (~63 mg, 79% yield), LC-MS (ESI) (m/z): calculated for $C_{37}H_{60}N_2O_{13}$: 740.84**. TMP-PEG$_{15}$-NH$_2$**: a sticky solid (~66 mg). LC-MS (ESI) (m/z): calculated for $C_{47}H_{80}N_2O_{18}$: 960.54, found 961.5 (M+1). **TMP-PEG$_{23}$-NH$_2$**: a sticky solid (~59 mg). LC-MS (ESI) (m/z): calculated for $C_{63}H_{112}N_2O_{26}$: 1312.75, found 1313.7 (M+1).

**Synthesis of TMP-PEG$_n$-NHCO-Biocytin-azide** (n = 1, 10, 15 and 23): To the solution of N-(azido PEG4) biocytin (10 mg, 0.15 mmol) in DMF was added diisopropyl ethylamine (22.59 uL, 0.13 mmol) then HATU (17 mg 0.46 mmol) sequentially. The reaction mixture was stirred for 20 minutes at room temperature. TMP-PEG$_n$-NH$_2$ (0.21 mmol, 20 mg) in DMF was added slowly to the above reaction mixture and allowed to stir at room temperature for 12 h. The solvent was removed under reduced pressure, and the residue was purified by silica gel column chromatography.

**TMP-PEG$_{10}$-NHCO-Biocytin-azide** eluted with chloroform, methanol and 28% ammonia (10:1:0.5), recovered as a light red pellet. (88 mg, 75.61% yield). LC-MS (ESI) (m/z): calculated for $C_{64}H_{105}N_9O_{21}S$: 1368.6, found 1368.7 (M+1).

**TMP-PEG$_{15}$-NHCO-Biocytin-azide** was eluted from the column using a mixture of chloroform, methanol and 28% ammonia solution (8:2:0.5) recovered as a light red pellet. LC-MS (ESI) (m/z): calculated for $C_{74}H_{125}N_9O_{26}S$: 1588.8, found 1589.8 (M+1).

**TMP-PEG$_{23}$-NHCO-Biocytin-azide** was eluted from the column using mixture of chloroform, methanol and 28% ammonia solution (8:2:0.5) recovered as a light red pellet. LC-MS (ESI) (m/z): calculated for $C_{90}H_{157}N_9O_{34}S$: 1941.3, found 1942.8 (M+1).

#### Functionalization of secondary antibody with DBCO-NHS ester

The secondary antibody (2mg/ml) was mixed with 50 times molar excess of DBCO-PEG13-NHS ester (a 50 mM stock solution in DMSO) then left at 4°C overnight on a rotator. The mixture was dialyzed (cut off 10 kDa) against 1X PBS (137 mM NaCl, 2.7 mM KCl, 10 mM $Na_2HPO_4$, and 1.8 mM $KH_2PO$, pH 7.4) at 4°C for 8 h, to remove the unreactive DBCO-PEG13-NHS.

#### Copper-free click reaction with TMP-PEG$_n$-NHCO-Biocytin-azide

The TMP-PEG$_n$-NHCO-Biocytin-azide pellets were dissolved in ethanol and a 1:50 dilution was prepared by dissolving TMP-PEG$_n$-NHCO-Biocytin-azide in nuclease-free water. The concentration of each compound was measured by UV spectroscopy, OD at 250 nm (extinction co-efficient at 25000). The antibody-DBCO was mixed with a TMP-$PEG_n$-NHCO-Biocytin-azide and then left at 4°C overnight on a rotator. The samples were dialyzed against 1X phosphate buffered saline overnight at 4°C and qualified by the western blot analysis shown in **Fig 2** [1, 2].

**Biological and Biochemical Materials and Methods**

**Cell Culture.** Hela cell lines were grown in DMEM media supplemented with 10% fetal bovine serum and Penicillin-streptomycin antibiotics (Gibco, USA).

Dig-TMP was synthesized as per the reported protocol [3].

**Table 1.** List of antibodies, their source, catalog and working dilutions

| Antibody | Source | Catalog | Dilutions |
|---|---|---|---|
| Rabbit anti Digoxigenin | Invitrogen | 710079 | 1: 200 |
| Mouse anti Digoxigenin | Abcam | Ab420 | 1:200 |
| HRP anti mouse | Invitrogen | 31432 | 1:1000 |
| Mouse anti-Biotin | Jackson Immunoresearch | 200-002-211 | 1:1000 |
| Anti-γ-H2A.X (phospho S139) antibody | Millipore sigma | 05-636 | 1:500 |
| Mouse anti PCNA | Abcam | AB29 | 1:100 |
| Rabbit anti MCM2 | Abcam | AB108935 | 1:100 |
| Biotinylated goat anti streptavidin | Vector labs | BA-0500 | 1:200 |
| QDot 655 anti-rabbit | Invitrogen | Q11421MP | 1:2500 |
| QDot 705 anti-mouse | Invitrogen | Q11062MP | 1:2500 |
| QDot 705 Streptavidin | Invitrogen | Q10161MP | 1:2500 |
| Alexa 488 anti-Rat | Invitrogen | A11006 | 1:200 |
| Alexa 488 anti-Rabbit | Invitrogen | A11034 | 1:200 |
| Alexa 488 anti-Mouse | Invitrogen | A11001 | 1:200 |
| Alexa 633 anti-Mouse | Invitrogen | A21052 | 1:200 |
| Alexa 633 anti-Rabbit | Invitrogen | A21070 | 1:200 |
| N-(Azido-PEG4)-biocytin | Broad Pharm | BP23286 | |
| DBCO- $PEG_{13}$ $NH_2$ ester | Click chemistry tools | 1015-2 | |
| $H_2N$-$PEG_{10}$-$CH_2CH_2NH_2$ | Chemprep Inc | 281704 | |
| $H_2N$-PEG15-$CH_2CH_2NH_2$ | Pure PEG | 233615 | |
| $H_2N$-PEG23-$CH_2CH_2NH_2$ | Pure PEG | 233623 | |
| Goat anti Mouse | Jackson Immunoresearch | 115-005-003 | |
| Goat anti Rabbit | Jackson Immunoresearch | 111-005-003 | |
| Mouse anti Bioitn | Rockland | 200-301-098 | 1:100 |
| Rabbit anti biotin | Cell signalling | 5597S | |
| TMP-Cl | Berry associates | PS-5000 | |

**TMP or Dig-TMP and UVA Treatments.** Cells were incubated with 1 μM Dig-TMP in media (with N-Acetyl Cysteine 10 mM) for 20 min and then irradiated with UVA 3 $J/m^2$ at 37°C.

Identification of optimal PEG linker lengths- DNA fiber assay

This procedure featured two uses of psoralen to covalently attach tags to cellular DNA. In the first application, the Dig antigen linked to psoralen was conjugated to cellular DNA in live cells. In the second the biotin tag in the Y conjugates was attached to the DNA in the fixed and permeabilized cells.

Cells were grown on 12 well plates to 40% confluency and then incubated with CldU (10 μM) for 12 hr. Then they were incubated with Dig- TMP in media for 20 min at 37°C and exposed to UVA at 3 $J/cm^2$ to covalently link the Dig tag to cellular DNA. The cells were washed with 1X PBS and incubated in methanol at -20°C for 20 min followed by treatment with 0.5 % Triton X-100 buffer (100 mM glycine, 0.2mg/ml EDTA, 1% BSA, 0.5% Triton X-100 in PBS) for 10 min at 4°C. They were incubated with 100 ug/ml RNAse for 30 min at 37°C, washed with PBS, and then blocked with 5% BSA, 10% goat serum in PBS for 1 h at room temperature. The blocking solution was removed followed by incubation with rabbit anti-Dig antibody for 1 hr at room temperature. After washing with 1X PBST and treat with respective Y conjugates for 1 h at room temperature or overnight at 4°C, after which they were washed with PBS-T to remove unbound compounds. They were then exposed to UVA at 3 $J/cm^2$ to covalently link the conjugates to cellular DNA. Fibers were spread and the Dig tag was identified by incubation with a mouse anti-Dig primary antibody and a quantum dot (655) linked to a goat anti-mouse secondary antibody. The biotin tag was displayed by incubation with a rabbit primary antibody against biotin and a goat anti-rabbit secondary conjugated to quantum dot (705).

**Western blot detection**

To assess the integrity of the biotin on synthesized psoralen-ADC's, conjugates were denatured by boiling at 95°C for 5 min in NuPAGE Sample Buffer (Invitrogen) containing β-mercaptoethanol (BME). Denatured samples were resolved on 12% Tris-glycine precast gels and transferred onto 0.2 μm PVDF membrane for 1 h at 4°C using standard procedure. After transfer membranes were washed twice with PBS, blocked for 1 h at room temperature in blocking buffer (5% nonfat dry milk in PBS-T with 0.1% Tween-20), 1 h with mouse anti-biotin (1:1000); 1 h in Horseradish peroxidase (HRP) anti mouse: (1:1000) at room temperature. During each step, membranes were washed three times with PBS-T for 10 min. After the incubation HRP secondary antibody proteins were detected using enhanced chemiluminescence reagents and visualized using Biorad Chemidoc system.

**Immunofluorescence**

HeLa cells were grown on Mattek glass-bottomed petri plates coated with Cell-Tak™ (Corning). Next day, cells were incubated with CSK-R buffer (10 mM PIPES pH 6.8, 100 mM NaCl, 300 mM Sucrose, 3 mM MgCl2, 0.7% Triton X-100, 0.3 mg/ml RNAse A) and fixed with 4% Formaldehyde in PBS at room temperature. After washing with PBS, cells were incubated in ice-cold methanol at -20°C for 20 min and treated with 0.5 % Triton X-100 buffer (100 mM glycine, 0.2mg/ml EDTA, 1% BSA, 0.5% Triton X-100) for 10 min at 4°C, followed by treatment with 100 ug/ml RNase A for

30 min at 37°C. Subsequently, cells were blocked (5% BSA, 10% Goat serum in PBS) for 1 h at room temperature and incubated with respective conjugated Y conjugates for 1 h at room temperature or overnight at 4°C followed by UVA or no UVA. After washing three times with PBS-T (0.1 % Tween-20) cells were incubated with Mouse anti-biotin (1:200) for 1 h at room temperature or overnight at 4°C, 1 h with Alexa 633 goat anti-Mouse (1:200) at room temperature. Finally, cells were treated with Prolong Gold with DAPI (pan nuclear DNA stain) and visualized using a Zeiss Axiovert 200M microscope at x63 magnification.
